# Cycloheximide resistant ribosomes reveal adaptive translation dynamics in *C. elegans*

**DOI:** 10.1101/2025.05.07.652686

**Authors:** Qiuxia Zhao, Blythe Bolton, Reed Rothe, Reiko Tachibana, Can Cenik, Elif Sarinay Cenik

## Abstract

Protein translation regulation is critical for cellular responses and development, yet how disruptions during the elongation stage shape these processes remains incompletely understood. Here, we identify and validate a single amino acid substitution (P55Q) in the ribosomal protein RPL-36A of *Caenorhabditis elegans* that confers complete resistance to high concentrations of the elongation inhibitor cycloheximide (CHX). Heterozygous animals carrying both wild-type RPL-36A and RPL-36A(P55Q) exhibit normal development but intermediate CHX resistance, indicating a partial dominant effect. Leveraging RPL-36A(P55Q) as a single-copy positive selection marker for CRISPR-based genome editing, we introduced targeted modifications into multiple ribosomal protein genes, confirming its broad utility for altering essential loci. In L4-stage heterozygotes, where CHX-sensitive and CHX-resistant ribosomes coexist, ribosome profiling revealed increased start-codon occupancy, suggesting early stalling of CHX sensitive ribosomes. Chronic CHX reduced ribosome collisions, evidenced by fewer disomes and unchanged codon distributions in monosomes. Surprisingly, prolonged elongation inhibition did not activate well characterized stress pathways–including ribosome quality control (RQC), the ribotoxic stress response (RSR), or the integrated stress response (ISR)–as indicated by absence of changes in RPS-10 ubiquitination, eIF2α phosphorylation, PMK-1 phosphorylation, or the transcriptional upregulation of ATF-4 target genes. Instead, RNA-normalized ribosome footprints revealed gene-specific changes in translation efficiency, with nucleolar and P granule components significantly decreased while oocyte development genes were increased. Consistent with these observations, we detected premature oogenesis in L4 animals, suggesting that partial translation elongation inhibition reshapes translation efficiency, to fine-tune developmental timing.

## INTRODUCTION

Translational regulation is integral to developmental transitions across multiple organisms, including *Drosophila*, *Xenopus* and zebrafish, where it controls critical processes such as the maternal-to-zygotic transition and the switch from stem cell self-renewal to differentiation (Weil et al. 2012; Lee et al. 2014; Teixeira and Lehmann 2019; Zhang et al. 2022). These regulatory mechanisms establish and maintain cell identity, tissue homeostasis, and tumor suppression (Buszczak et al. 2014). In *Drosophila*, for instance, translational control during axis specification involves remodeling of transport ribonucleoprotein (RNP) complexes and the dynamic partitioning of mRNAs between the translationally active edge and the silent core of P bodies, a process necessary for establishing proper embryonic patterning (Weil et al. 2012). Similarly, in *Caenorhabditis elegans*, the translational regulators LIN-41 and OMA-1/2 coordinate oocyte growth and the spatiotemporal regulation of meiotic maturation (Spike et al. 2014).

Among the stages of protein synthesis, elongation is both the longest and the most energy intensive, making it particularly susceptible to defects. Disruptions in elongation can lead to translational infidelity and ribosome stalling, which have been linked to neurodegeneration, cancer, and cardiovascular diseases (Liu and Proud 2016; Kapur et al. 2017; Wang and Sun 2023). Beyond these pathological contexts, elongation also allows cells to respond to stress through ribosome pausing and collisions, events that trigger ribosome quality control (RQC), the integrated stress response (ISR), and the ribotoxic stress response (RSR), thereby maintaining cellular homeostasis (Simms et al. 2017; Vind, Snieckute, et al. 2020; Wu et al. 2020; Barros et al. 2023).

Investigating how ribosome collisions influence growth and differentiation in multicellular organisms remains challenging. *C. elegans* offers a powerful system to address this gap, thanks to its invariant cell lineage, transparency, and genetic tractability. These features enable both tissue-specific and organism-wide analyses of translation regulation during development.

A common method for studying translation elongation involved the elongation inhibitor cycloheximide (CHX), which blocks the E site on the large subunit (Schneider-Poetsch et al. 2010; Garreau De Loubresse et al. 2014; Santos et al. 2019). CHX has been invaluable in dissecting ribosome function across eukaryotes (Mohammad et al. 2019; Wu et al. 2019; Wu et al. 2020). Intriguingly, certain organisms exhibit inherent CHX resistance. For example, *Kluyveromyces lactis* harbors a Pro-to-Gln substitution in RPL41 that confers resistance; a variant also found in *Candida maltosa* and generated in *Saccharomyces cerevisiae* (Takagi et al. 1992; Dehoux et al. 1993). In yeast, coexisting CHX-sensitive and CHX-resistant ribosomes have been shown to promote ribosome collisions and activate the RQC, offering a valuable model for investigating collision mediated stress (Simms et al. 2017).

Although ribosome associated stress responses are well-characterized in yeast and mammalian cell cultures (Meydan and Guydosh 2020; Vind, Genzor, et al. 2020; Yan and Zaher 2021; De and Mühlemann 2022), their functional impact on development in multicellular animals remain less understood. *C. elegans,* which shares many of the same translation regulatory mechanisms with higher eukaryotes (Rhoads et al. 2006; Mehta et al. 2010; Blazie et al. 2021), provides a unique model to dissect how collisions, and subsequent stress responses regulate gene expression and developmental progression at the organismal level.

Here, we investigate the consequences of inhibiting translation elongation on *C. elegans* development. Specifically, we identify *rpl-36A*, encoding the ribosomal protein RPL-36A, as the *C. elegans* homologue of *K. lactis* L41 and show that a P55Q substitution confers strong CHX resistance in vivo. We then utilize RPL-36A(P55Q) as a positive selection marker for CRISPR genome editing to introduce targeted modifications in ribosomal protein genes, including a single amino acid substitution (*rps-23(R67K)*) and the addition of a C-terminal *HA::AVI::TEV* tag to the *rpl-16*. By examining *C. elegans* heterozygotes carrying both CHX-sensitive (RPL-36A) and CHX-resistant (RPL-36A(P55Q)) ribosomes, we observe global changes in ribosome occupancy, most notably increased monosome accumulation at start codons and reduced disome occupancy across coding regions, indicative of altered translation dynamics under partial elongation inhibition. Integrating RNA-seq and Ribo-seq revealed that these shift predominantly genes involved in gametogenesis, causing oogenesis to initiate before spermatogenesis. Moreover, translational regulators, including OMA-1, OMA-2, and LIN-41, are overexpressed under CHX stress, with increased ribosome occupancy on their downstream targets. Together, our findings suggest that moderate chronic inhibition of translation elongation disrupts the precise coordination required for sperm and oocyte development in *C. elegans*. By defining how ribosome collisions and translational stress responses shape gene expression and cell-fate decisions, this provides new insights into the role of translational elongation regulation in multicellular development.

## RESULTS

### RPL-36A(P55Q) confers cycloheximide resistance in *C. elegans*

In *Kluyveromyces lactis,* a glutamine residue (Q56) in the large subunit ribosomal protein L41 confers resistance to high concentrations of cycloheximide (CHX) (Dehoux et al. 1993). To determine whether this residue and its associated function are conserved, we compared homologous sequences from *Saccharomyces cerevisiae*, *Mus musculus*, *Homo sapiens*, *Caenorhabditis elegans,* and *Drosophila melanogaster*. A proline in position 56 (P56) is present in all examined species except *K. lactis* (**Fig 1A**), whose L41 homologs nonetheless share high overall sequence identity with their counterparts (**Fig S1A**, **S1B**). In *C. elegans,* RPL-36A is homologous to *K. lactis* L41, carrying proline at residue 55 instead of glutamine (**Fig 1A, 1B)**.

**Figure 1.**
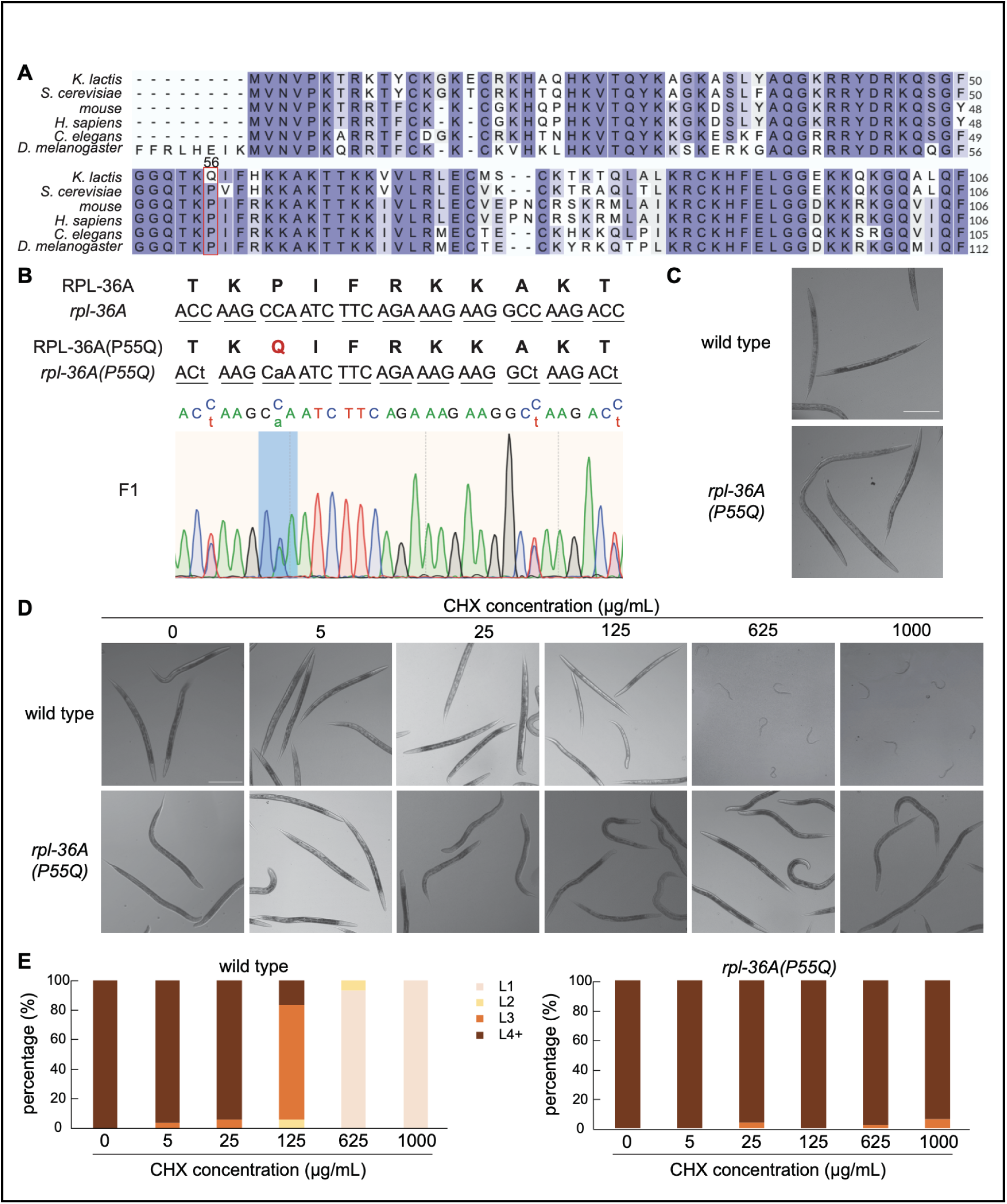
*rpl-36A(P55Q)* mutants are resistant to high concentrations of CHX. **(A)** The amino acid sequence alignments of CHX resistant *K. lactis* L41 orthologues in *S. cerevisiae* (RPL42A), mouse (RPL36A), *H. sapiens* (RPL36A), *C. elegans* (RPL-36A), and *D. melanogaster* (RPL36A) show that Q56 is unique to *K. lactis* L41. **(B)** A genetic mutation at residue 55 of RPL-36A, substituting proline with glutamine, was introduced in *C. elegans* using CRISPR-Cas9. **(C)** *rpl-36A(P55Q)* animals display a wild-type phenotype. **(D)** Synchronized embryos of wild type and *rpl-36A(P55Q)* were treated with different concentrations of CHX for 2 days at 23 ℃. Scale bar in **C** and **D**, 250 µm. **(E)** Quantification of larval stages in wild type (left) and *rpl-36A(P55Q)* animals (right) under different concentrations of CHX. N>40.

To assess whether substituting P55 with Q confers CHX resistance in *C. elegans*, we used CRISPR-Cas9 to introduce P55Q mutation at the endogenous *rpl-36A locus* (**Fig 1B**). Homozygous *rpl-36A(P55Q)* animals were viable and phenotypically indistinguishable from wild-type animals (**Fig 1C**), indicating that RPL-36A(P55Q) does not compromise basic ribosome function.

Next, we examined post embryonic development under various CHX concentrations (0, 5, 25, 125, 625, 1000 µg/ml) at 23 ℃ for 2 days. Both wild-type and *rpl-36A(P55Q)* animals developed to the L4 or young adult stage without CHX (**Fig 1D, 1E**). However, wild-type animals displayed dose-dependent growth inhibition and eventually arrested at the L1 stage at CHX concentrations equal to or higher than 625 µg/ml (**Fig 1D, 1E**), in line with prior work linking limited ribosomal function to development arrest (Dalton and Curran 2018; Cenik et al. 2019; Zhao et al. 2023). In contrast, *rpl-36A(P55Q)* animals continued normal development even at the highest concentrations tested (625 or 1000 µg/ml), confirming that RPL-36A(P55Q) confers robust CHX resistance in *C. elegans* (**Fig 1D, 1E)**.

### Utilizing RPL-36A(P55Q) as a CHX-resistance marker for CRISPR co-conversion

Standard CRISPR-Cas9 genome editing in *C. elegans* often yields heterozygous modifications because injection occurs near the hermaphrodite gonad, rather than the spermatheca (Arribere et al. 2014). Typical co-conversion strategies use visible phenotypic markers (*dpy-10* or *sqt-1*), to track desired edits at an unlinked locus (Arribere et al. 2014). CHX-based selection can streamline this process by reducing phenotypic scoring, and increasing selection efficiency.

To determine whether *rpl-36A(P55Q)* can serve as a dominant selection marker, we crossed *rpl-36A(P55Q)* mutants with a balancer strain (*mIn1*), which carries a large inversion that suppresses recombination over a significant portion of chromosome II. The balancer is marked by two phenotypic features: *dpy-10*, which causes a Dumpy body shape when homozygous, and a *myo-2::GFP* transgene that drives fluorescence in the pharyngeal muscles (Edgley and Riddle 2001). In the absence of CHX, *rpl-36A(P55Q)*/*mIn1* heterozygotes were phenotypically similar to *rpl-36A(P55Q)* homozygotes, and *mIn1/mIn1* homozygotes displayed a characteristic Dumpy body shape (**Fig 2A**). Upon exposure to 1 mg/ml CHX for 2 days, *rpl-36A(P55Q)* homozygous animals reached adulthood, whereas *mIn1/mIn1* homozygotes were arrested at the L1 stage (**Fig 2A**). *rpl-36A(P55Q)*/*mIn1* heterozygotes grew to the L3-L4 stage, indicating a dose-dependent, dominant form of CHX resistance (**Fig 2A**). Notably, *rpl-36A(P55Q)* progeny from heterozygous parents were smaller under CHX than those from *rpl-36A(P55Q)* parents, likely reflecting maternally deposited CHX-sensitive ribosomes that persist through postembryonic development (Cenik et al. 2019).

**Figure 2.**
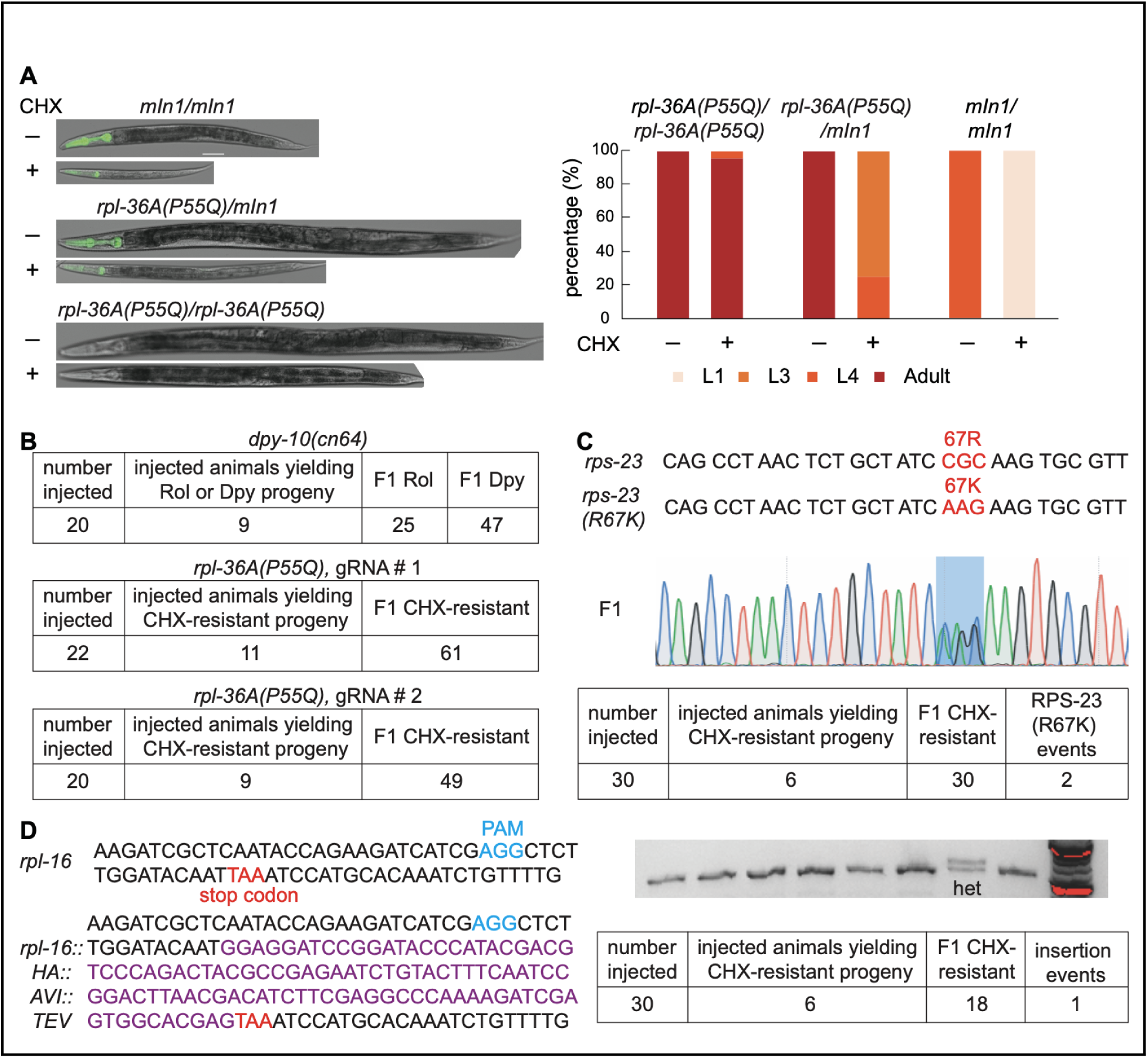
Utilizing the single-copy CHX-resistant *rpl-36A(P55Q)* gene as an efficient CRISPR co-conversion marker. **(A)** Synchronized *rpl-36A(P55Q)/mIn1* progeny were grown on 1 mg/ml CHX plates at 23 ℃ for 2 days. Scale bar: 50 µm. Quantification of larval stages of animals, n=37. (**B**) Injection efficiency for *dpy-10(cn64)*, *rpl-36A(P55Q)*. Twenty animals were injected for *dpy-10(cn64).* Two gRNAs targeting *rpl-36A(P55Q)* were used in separate injection experiments. See main text for description of Dpy, Rol and CHX-resistant phenotypes. (**C**) Scheme of the *rps-23(R67K)* locus. The R67K mutation (Arg→Lys) is highlighted in red. Sanger sequencing of the single-worm PCR results from an F1 CHX-resistant individual and the R67K event is shown in blue shading. (**D**) Schematic of the *rpl-16(rpl-16::HA::AVI::TEV)* locus. A fragment (HA::AVI::TEV) is inserted at the C-terminus of rpl-16 locus, just before the stop codon (TAA). The PAM sequence is indicated in blue. Single-worm PCR was performed to detect the insertion in F1 CHX-resistant progeny.

The co-conversion strategy is widely used in CRISPR/Cas9 genome editing, wherein a dominant phenotypic oligonucleotide-templated conversion event at one locus is used to enrich for desired modifications at another unlinked locus (Arribere et al. 2014). Mutations in *dpy-10* and *sqt-1* are commonly employed as effective co-conversion markers; however, they require manual selection of animals with the dominant phenotype (Arribere et al. 2014). Building on our observation that the *rpl-36A(P55Q)* allele confers a dominant, dose-dependent resistance to CHX, we tested whether this allele could serve as a co-CRISPR selection marker to facilitate the generation of small nucleotide edits. F1 progeny were then screened for CHX resistance.

Before evaluating *rpl-36A(P55Q)* as a conversion marker, we compared its editing efficiency to that of *dpy-10(cn64)*. Two gRNAs targeting *rpl-36A* were designed and used in CRISPR /cas9 injections. For CHX-based selection, 500 µl of 10 mg/ml CHX was applied to the surface of each 10 ml NGM plate on the second day post-injection, achieving a final concentration of approximately 500 µg/ml CHX. For *rpl-36A(P55Q)*, gRNA #1 produced CHX-resistant F1s in 11 out of 22 injected animals (61 total F1s), and gRNA #2 yielded resistant F1s in 9 out of 20 injected animals (49 total F1s) (**Fig 2B**). In comparison, *dpy-10(cn64)* yielded roller or dumpy F1s in 9 out of 20 injected animals, with 72 phenotypically marked progeny in total (**Fig 2B**), demonstrating that *rpl-36A(P55Q)* achieves comparable genome editing efficiency to *dpy-10*.

Leveraging this CHX resistance, we introduced two edits into ribosomal protein genes: a single amino acid substitution in *rps-23(R67K)* (**Fig 2C**), and a C-terminal HA::AVI::TEV tag in *rpl-16* (**Fig 2D**). Following CRISPR injections, we applied 500 µl of 10 mg/ml CHX onto the surface of each 10 ml NGM plate with animals, achieving a final concentration ∼500 µg/ml CHX. We then screened F1 progeny for CHX resistance, performed single worm PCR, and confirmed the edits by sequencing. Among 30 injected worms, we obtained two *rps-23(R67K)* lines out of 30 CHX-resistant F1 progeny, and one *rpl-16(rpl-16::HA::AVI::TEV*) line out of 18 CHX-resistant F1 progeny. Thus, RPL-36A(P55Q) provides an effective, single-copy selection marker for CRISPR editing of essential genes.

### Ribosome stalling at start codons and reduced disome occupancy under chronic translation elongation inhibition

Leveraging heterogeneity in ribosome composition to detect ribosome stalling events under elongation stress was inspired by prior studies in yeast and cell culture (Ref). *rpl-36A(P55Q)/+* heterozygotes displayed an intermediate level of CHX resistance, prompting us to model these conditions in vivo and directly assess the consequences of ribosome collisions.

To determine how a mixture of CHX-resistant (RPL-36A(P55Q)) and CHX-sensitive (wild-type) ribosomes affects translation elongation, we performed Ribo-seq, Diso-seq, and RNA-seq on L4 stage *rpl-36A(P55Q)/+* heterozygotes, treated with and without CHX. All datasets originated from the same biological replicates, and showed high reproducibility (**Fig S2A,** Spearman coefficient ∼ 0.95 across biological replicates).

Ribosome-protected fragments (RPFs) were predominantly 29-33 nucleotides, while disome-protected fragments (DPFs) were 62-68 nucleotides (**Fig 3A**). Both monosome and disome protected fragments mapped mainly to coding regions and exhibited 3-nt periodicity (**Fig 3B, 3C, Fig S2B**). Under CHX treatment, monosome footprints accumulated at the start codon and decreased near the stop codon (**Fig 3B**), suggesting early stalling in CHX-sensitive ribosomes (Lee et al. 2012). Disome footprints also showed enrichment at the start codon, and were reduced across the coding regions and the stop codon (**Fig 3C, D**).

**Figure 3.**
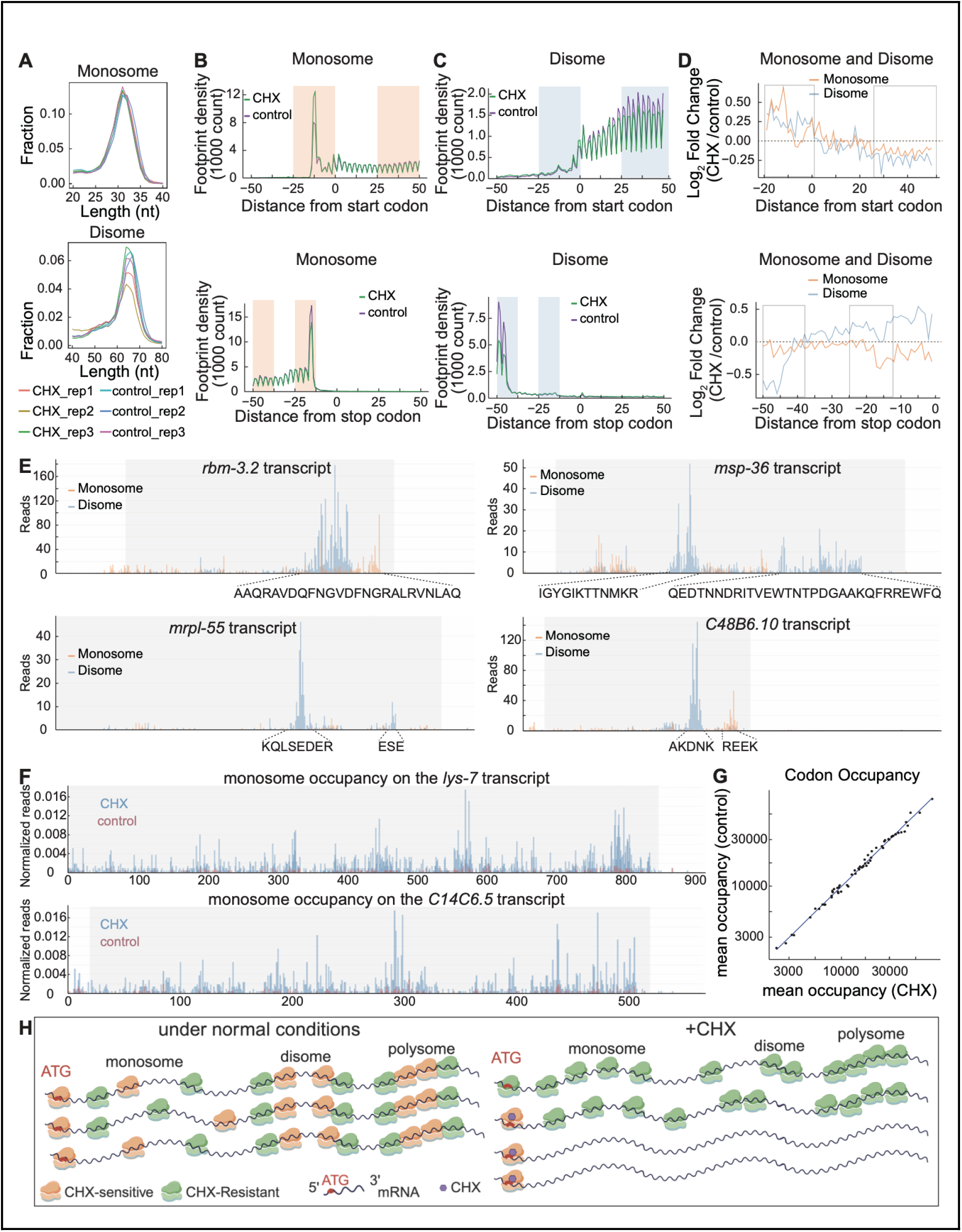
Ribosome occupancy changes in response to translation elongation inhibition. **(A)** Length distribution of ribosome-protected fragments (RPFs) (top) and disome-protected fragments (DPFs) (bottom) for *RPL-36A(P55Q)/+* animals under CHX (625 µg/ml) treatment and their controls. **(B)** Meta-gene analysis of monosome occupancy aligned at the start and stop codons in heterozygous animals with or without CHX treatment. **(C)** Meta-gene analysis of disome occupancy aligned at the start and stop codons in heterozygous animals with or without CHX treatment. The rectangles in (**B**) and (**C**) highlight the same positions to facilitate comparisons across monosome and disome occupancy. **(D)** Log_2_ fold changes in monosome and disome occupancy aligned at the start and stop codons in the heterozygous animals with or without CHX. **(E)** Distribution of monosome and disome occupancy across coding regions under normal conditions. Representative examples of four genes (*rbm-3.2*, *msp-36*, *mrpl-55* and *C48B6.10*) show enriched disome footprints and corresponding amino acid sequences. **(F)** Distribution of monosome occupancy across coding regions under CHX treatment and control conditions. Representative examples of two genes (*lys-7* and *C14C6.5*) show a significant increase in monosome occupancy in CHX-treated samples compared to controls. The Y-axis represents the normalized value per 1,000 total reads. In **E** and **F**, the light gray boxes along the X-axis indicate coding regions, while the white background represents the 5’UTR and 3’UTR, respectively. **(G)** Scatter plot comparing codon-specific ribosome occupancies in heterozygous animals treated with or without CHX. **(H)** Model illustrating translation elongation inhibition in CHX-treated animals with mixed CHX-sensitive or CHX-resistant ribosomes. Under normal conditions, heterozygous animals efficiently translate mRNAs into proteins. Upon CHX treatment, sensitive ribosomes stalled at the start codon region, leading to reduced ribosomes progressing along the mRNA and fewer ribosome collisions. BioRender.

Consistent with prior observations (Zhao et al. 2021), genes with higher monosome occupancy tended to have more disomes (**Fig S2C, Table S1,** rho =0.93, p-value < 2.2e-16 control; rho = 0.88, p-value < 2.2e-16, CHX, Spearman rank correlation of average DPFs and RPFs divided by average RNA levels). Despite this trend, some genes showed disproportionately higher disome occupancy relative to monosome occupancy, suggesting they are particularly prone to ribosome collisions (**Fig S2C, Table S1**). To explore this further, we examined ribosome occupancy profiles on transcripts that showed a disome-to-monosome ratio ≥ 2.7 with at least 100 disome reads under normal conditions. Only 12 genes met these criteria among all those analyzed (**Table S2**): *col-12, msp-36, ctc-3, ctc-1, ctc-2, nduo-3, rmb-3.2, nlp-34, C48B6.10, Y48G8AL.12, mrpl-55,* and *dib-1*. We visualized disome and monosome positioning on four transcripts (**Fig 3E**) (Chacko et al. 2024). In these examples (*rmb-3.2, msp-36, mrpl-55 and C48B6.10*), disome footprints generally accumulated near the 3’ end, often immediately downstream of monosome stall sites. Although we observed no universal amino acid bias across disome-enriched regions, disome peaks spanned diverse sequence contexts. For example, in *mrpl-55* and *C48B6.10*, we detected clusters of charged residues (e.g., lysine, glutamic acid, aspartic acid, arginine), whereas *rbm-3.2* and *msp-36* contained a mix of polar, hydrophobic and acidic residues under the same peaks (**Fig 3E**). These results suggest that stalling could be influenced by codon usage or by the presence of acidic residues in the nascent peptide channel (Arpat et al. 2020; Zhao et al. 2021), affecting ribosome elongation.

We further investigated the relationship between ribosome stalling and amino acid residues in untreated worms by translating the monosome and disome read sequences into peptide sequences. The percentage of each amino acid in the total disome or monosome sequence compared to the percentage in the CDS revealed little to no differences for most amino acids (**Fig S3A**). However, both monosome- and disome-occupancy exhibited higher mean sequence percentage of lysine, which has been associated with ribosome collisions in the case of polylysine tracts in yeast (Zhao et al. 2021). To identify longer sequence elements related to ribosome stalling, motif enrichment analysis of translated disome sequences was performed using STREME (Bailey 2021). The most highly enriched motif contained a tri-residue repeat (**Fig S3B, Table S3**). This pattern indicates the collagen repeat G-X-Y where X and Y are most commonly proline and hydroxyproline (Ramachandran 1956). Within the STREME test set of sequences, the enriched collagen motif manifested in at least 50 disome reads across 36 genes, all of which encode collagen peptides or predicted structural constituents of the cuticle (**Fig S3C**). The gene with the greatest number of reads containing the collagen motif was *col-77*, and when monosome and disome coverage were plotted, a distinct disome peak corresponding to the motif was observed (**Fig S3D**). The sharp disome peak associated with the collagen motif suggests a relationship between ribosome stalling and the helical structure of the collagen tri-residue repeat. This may relate to the transition from an unstructured peptide region to the structured G-X-Y repeat since the progression through structured-unstructured-structured peptide configurations has demonstrated disome enrichment (Arpat et al. 2020).

To evaluate whether translation elongation inhibition alters gene-specific translation efficiency in a codon-or sequence-dependent manner, we examined the monosome occupancy in genes that exhibited at least a 2-fold increase in monosome occupancy in response to CHX (**Table S4**). Among these, the six representative genes (*lys-7, C14C6.5, irg-5, ule-3, D1054.10,* and *perm-2*) showed the highest fold-changes in monosome occupancy in CHX-treated animals compared to the control. However, we found no localized enrichment of ribosome accumulation within the coding regions (**Fig 3F, S4**). Furthermore, codon occupancy analysis revealed no CHX-induced preference under monosome peaks, regardless of treatment (**Fig 3G**). These observations suggest that in CHX-treated animals, which harbor both CHX-sensitive and CHX-resistant ribosomes, elongation stalls predominantly at early stages, preventing most ribosomes from progressing through the coding sequence. Nevertheless, a compensatory mechanism may limit ribosome collisions, enabling sufficient protein synthesis to maintain cellular function despite impaired elongation (**Fig 3H**).

### ISR, RQC, and RSR signaling pathways are not activated in response to chronic translation elongation inhibition

Organisms rely on stress response pathways to maintain homeostasis and ensure survival under ribosome collisions. Given the decrease in disome occupancy observed in CHX-treated animals (**Fig 3C**), we investigated whether the downstream stress response pathways were induced. The major surveillance mechanisms pathways for resolving ribosome stalling include the ribosome-associated quality control (RQC) pathway, the integrated stress response (ISR), and the ribotoxic stress response (RSR) (Meydan and Guydosh 2020; Vind, Genzor, et al. 2020; Yan and Zaher 2021; De and Mühlemann 2022).

In heterozygous animals, CHX inhibits translation elongation by targeting sensitive ribosomes, causing ribosome stalling. Similarly, wild-type animals treated with intermediate CHX concentrations exhibit partial elongation inhibition (Simms et al. 2017). To determine if these stress pathways were activated, we systematically examined their key components in wild-type animals under intermediate CHX concentrations.

The RQC pathway in *C. elegans* requires ubiquitination of ribosomal proteins RPS-10 and RPS-20 (Monem et al. 2023). Wild-type animals treated with 125 µg/ml CHX spanned a range of developmental stages, while higher concentrations (625 µg/ml) caused arrest (**Fig 1E**). To assess RQC activation, we used a strain bearing endogenously HA-tagged *rps-10* exposed to intermediate concentrations of CHX (125, 250, and 375 µg/ml), and compared ubiquitinated RPS-10 (ubi-RPS-10) to total RPS-10 (**Fig 4A**). Western blots showed no increase in ubi-RPS-10 at these concentrations, indicating that moderate elongation inhibition via CHX does not robustly activate RQC (**Fig 4A**). Given that CHX treatment reduces ribosome density on coding sequences (**Fig 3C**), less RPS-10 may be available for ubiquitination overall.

**Figure 4.**
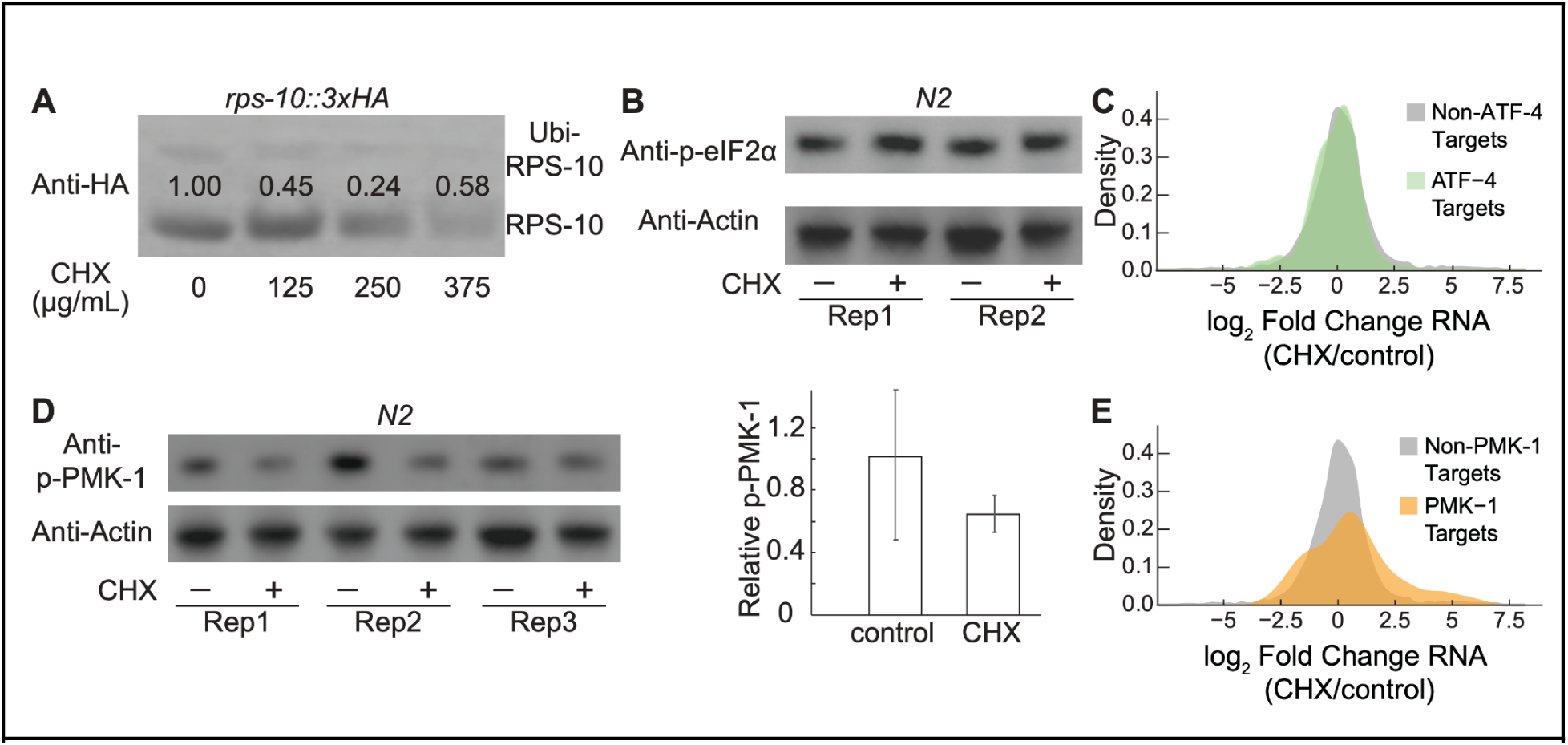
ISR, RQC, and RSR signaling pathways were not activated in response to chronic translation elongation inhibition. **(A)** An endogenously tagged *rps-10::3XHA* strain was used to assess the effects of different CHX concentrations (0, 125, 250, 375 µg/ml) on ribosome collision induced by translation elongation inhibition. Western blot analysis using an anti-HA antibody was performed to examine ubiquitylated RPS-10 (ubi-RPS-10) and total RPS-10 levels. Protein quantification was conducted using Fiji software, with the numbers above the gel lanes representing the relative ubi-RPS-10 levels normalized to RPS-10. **(B)** Wild-type animals were treated with or without 250 µg/ml CHX for three and two days, respectively. Western blot analysis was performed using antibodies against phosphorylated eIF2α (p-eIF2α) and Actin. **(C)** Log_2_ fold change (CHX/control) of ATF-4 target genes (green) and non-ATF-4 target genes (grey) were represented in a density plot in CHX-treated heterozygotes compared to controls, illustrating no significant upregulation of ATF-4 target genes in response to CHX treatment. **(D)** Wild-type animals were treated with or without 250 µg/ml CHX for three days and two days, respectively. Western blot analysis was performed using antibodies against phosphorylated PMK-1 (p-PMK-1) and Actin. Protein quantification was conducted using Fiji software, with p-PMK-1 levels normalized to Actin. Statistical analysis using a Student’s t-test showed no significant difference (p = 0.381). **(E)** Log_2_ fold change of PMK-1 target genes (orange) and non-PMK-1 target genes (grey) in CHX-treated heterozygotes compared to controls is represented in a density plot.

The ISR pathway is typically triggered by various stressors, such as nutrient deprivation and endoplasmic reticulum stress, leading to global suppression of protein synthesis via phosphorylation of eIF2α, and selective translation of the transcription factor ATF-4 (Ye et al. 2010; Horn et al. 2020; Statzer et al. 2022). To test whether CHX activates ISR, we examined phosphorylated eIF2α (p-eIF2α) via western Blot in wild-type animals treated with intermediate CHX concentrations. We found no significant increase in p-eIF2α in CHX-treated samples relative to controls (**Fig 4B**, mean CHX/control: 1.24 ± 0.30). Furthermore, as a proxy for ATF-4 activation, we examined the expression patterns of genes upregulated in response to ATF-4 overexpression (Statzer et al. 2022). RNA-seq analysis revealed no significant increase in the expression of these ATF-4 target genes in CHX-treated heterozygotes (**Fig 4C, S5A).** Instead, the mean log_2_ fold change for ATF-4 targets was significantly lower (mean= ∼-0.11) than that of other genes (mean = ∼0.1, t = −3.0611, p-value = 0.003, Welch two sample t-test). These results indicate that ISR was not activated in response to long-term CHX-induced ribosome stalling.

The ribotoxic stress response (RSR) is mediated by the MAP3K ZAKα, which senses ribosome stalling and activates p38 and JNK kinases (Snieckute et al. 2022). Although CHX-induced acute stress is a poor inducer of RSR activation (Vind, Snieckute, et al. 2020), we hypothesized that chronic stress might trigger this pathway. We examined the phosphorylated PMK-1 (p-PMK-1), the *C. elegans P38* MAPK ortholog, and no detectable increase in phosphorylated PMK-1 (p-PMK-1) was observed in CHX-treated animals (**Fig 4D**, mean CHX/control: 0.64 ± 0.37, p = 0.381). PMK-1 functions through a complex transcriptional program that supports tissue homeostasis (Yuan et al. 2023). We therefore analyzed the expression levels of genes that are differentially expressed in response to the loss of *pmk-1* (Yuan et al. 2023) in CHX-treated heterozygotes and controls. While RNA-seq indicated that PMK-1 responsive genes were modestly overexpressed (Welch two sample t-test, t = 2.43, p-value = 0.018; mean log_2_FC ∼0.62 for PMK-1 targets vs. ∼0.09 for all other genes, **Fig 4E, S5B**), the lack of increased p-PMK-1 and the small changes in PMK-1 targets suggest that the RSR pathway was not robustly activated.

Taken together, these results indicate that the RQC, ISR, and RSR signaling pathways are not strongly induced under chronic elongation inhibition. Our data support a model in which CHX-sensitive ribosomes stall near start codons, reducing both overall ribosome progression and potential collisions. Consequently, cells maintain protein synthesis and preserve function despite impaired elongation (**Fig 3H**).

### Activation of premature oocyte pre-maturation under chronic translation inhibition

To investigate how translation elongation inhibition affects global gene expression and ribosome dynamics, we performed RNA-seq and Ribo-seq analyses. Overall, we identified 2,220 genes with significantly altered mRNA levels (FDR < 0.05), 3,651 genes with significant changes in ribosome occupancy, and 209 showing significant differential ribosome occupancy relative to RNA abundance (**Fig 5A**, **Table 1, Table S5**). These genes exhibited distinct functional enrichments. Among the differentially expressed genes, those associated with double-strand break repair, regulation of meiotic cell cycle, and cell division were overexpressed at the mRNA level in CHX-treated animals (**Fig S6A, Table S6**). Similarly, DNA damage response and repair genes were also overexpressed in animals with ribosome biogenesis inhibition (Zhao et al. 2023). Although UV irradiation has been shown to activate the ribotoxic stress response (RSR) (Sinha et al. 2024), we did not detect a robust RSR activation here. Instead, we propose that the overexpression of double-strand break repair genes may reflect translation-associated stress or a broader response to ribosome stalling.

**Figure 5.**
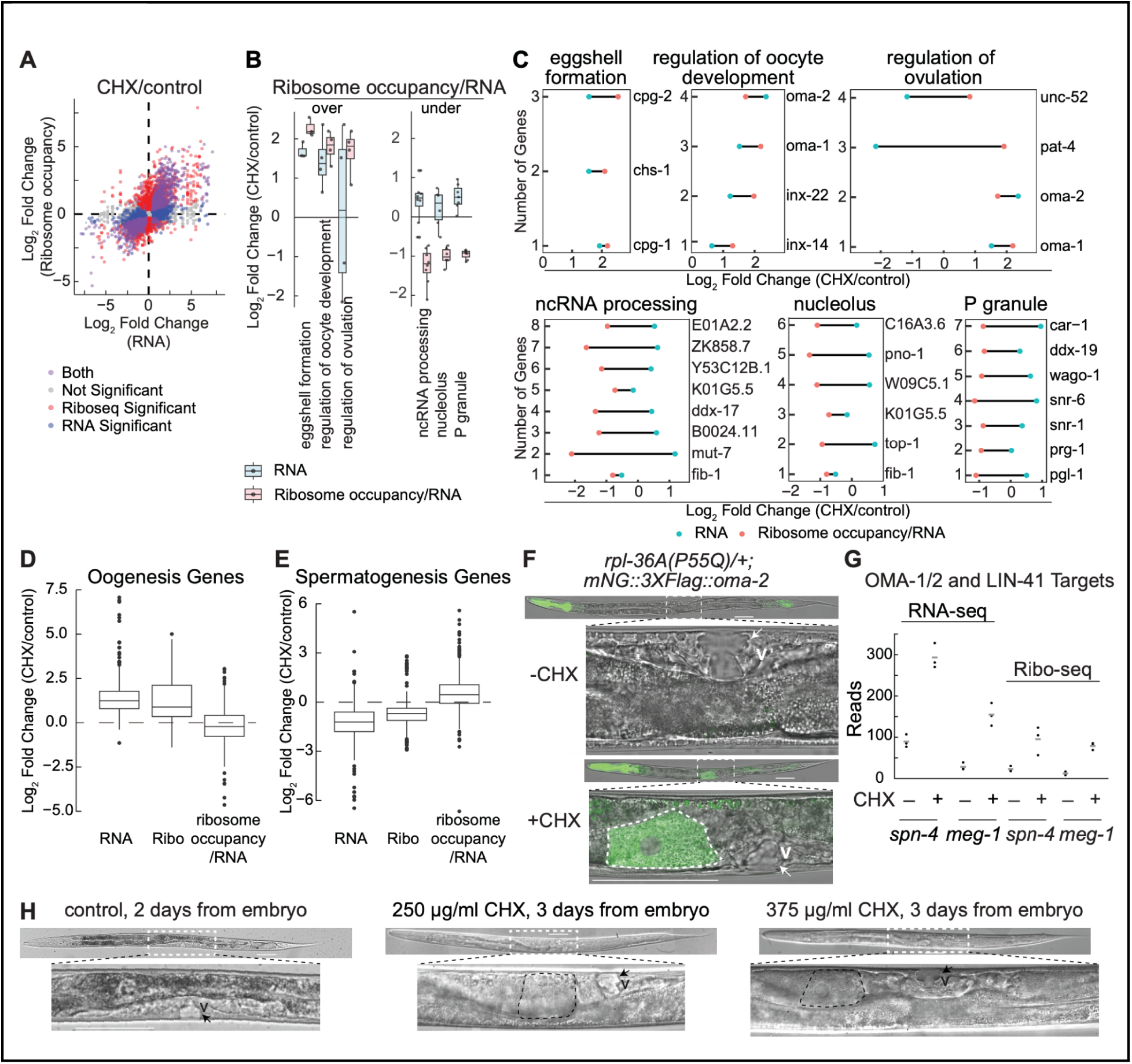
Differentially expressed genes in *rpl-36A(P55Q)/+* heterozygotes. **(A)** Log_2_ fold changes of genes predicted by RNA-seq (x-axis) were plotted against predicted ribosome occupancy (y-axis) in CHX-treated animals compared to control. Genes with significant changes at RNA level are shown in blue, those with significant changes of ribosome occupancy are shown pink, genes significantly changed at both levels are shown in purple, and genes that were not significant at either level are shown in gray. **(B)** Genes were characterized based on log_2_ fold change estimates from RNA-seq and ribosome occupancy/RNA. Significant gene ontology (GO) enrichment analyses were performed for each category: RNA and ribosome occupancy/RNA overexpression (left) and RNA overexpression with ribosome occupancy/RNA underexpression (right). **(C)** Gene annotation (GO) enrichment analyses for the categories shown in **(B)** were performed. The log_2_ fold change of RNA (blue) and ribosome occupancy/RNA (pink) for genes in six unique GO categories were plotted in response to CHX in the heterozygotes. **(D)** Log_2_ fold change values for oogenesis genes (Reinke et al. 2004; Ortiz et al. 2014) at RNA, Ribo, and ribosome occupancy/RNA levels (**Table S5**) were plotted in response to CHX treatment. **(E)** Log_2_ fold-change values of spermatogenesis genes at RNA, Ribo, and ribosome occupancy/RNA levels were plotted in response to CHX treatment. **(F)** Vulval phenotypes of L4-stage heterozygotes treated with or without CHX were analyzed. The *mNG::3XFLAG::oma-2* strain was crossed to *rpl-36A(P55Q)/+* heterozygotes. Under normal conditions, heterozygous animals displayed a wild-type phenotype. However, under CHX treatment, the animals were smaller than stage-matched controls, and their oocytes were premature despite displaying the classic L4-stage vulva. **(G)** Gene expression levels of OMA-1 and LIN-41 targets at the RNA and ribosome occupancy levels in heterozygotes under CHX stress. The y-axis represents the reads counts. **(H)** Wild-type animals treated with intermediate concentrations (250 µg/ml, 375 µg/ml) of CHX exhibited a premature oocyte phenotype. Arrows indicate the vulval structure. v: the vulvae. Scale bar: 50 µm.

**Table 1.** Numbers of differentially expressed genes at RNA, ribosome occupancy, and ribosome occupancy/RNA levels in heterozygotes under CHX treatment.

|  | RNA | Ribosome occupancy | Ribosome occupancy/RNA |
| --- | --- | --- | --- |
| Downregulated | 991 | 1763 | 118 |
| Upregulated | 1229 | 1888 | 91 |
| Not significant | 7833 | 6403 | 9844 |

In contrast, genes involved in fatty acid beta-oxidation and male meiosis chromosome segregation were underexpressed at the mRNA level, but showed increased ribosome occupancy relative to mRNA abundance (**Fig S6B**). This suggests a post-transcriptional compensatory mechanism in which reduced transcript availability is offset by enhanced translation efficiency.

Interestingly, genes related to eggshell formation, oocyte development, and ovulation regulation were overexpressed at the RNA level and showed increased ribosome occupancy in CHX-treated heterozygotes (**Fig 5B, 5C, Table S6**). Because L4-stage animals typically do not activate oogenesis or eggshell related genes until adulthood (Reinke et al. 2000), their elevated expression suggests premature initiation of the reproductive pathway. In contrast, functional categories with decreased ribosome occupancy per RNA included ncRNA processing, the nucleolus, and P granules (**Fig 5B, 5C, Table S6**), indicate a shift in translational priorities toward oogenesis at the expense of RNA metabolism and stress response pathways.

To investigate how CHX-induced ribosome stalling might influence germline cell-fate decisions, we focused on genes associated with oogenesis and spermatogenesis (Reinke et al. 2004; Ortiz et al. 2014). Oogenesis-related genes were overexpressed both at the RNA level and ribosome occupancy (**Fig 5D, Table S5**). In contrast, spermatogenesis-related genes were generally underexpressed and had reduced ribosome occupancy levels (**Fig 5E, Table S5**). These results imply that chronic elongation inhibition drives a bias toward premature oogenesis before activation of spermatogenesis-related genes.

To further characterize this shift, we examined the expression of *oma-1* and *oma-2*, which are primarily expressed in adult oocytes and function redundantly in oocyte maturation (Detwiler et al. 2001). Both genes were overexpressed at the RNA level and showed increased ribosome occupancy per RNA in CHX-treated heterozygotes (**Fig 5C, S6C**). When we used an mNG-tagged OMA-2 (Dickinson et al. 2017), we observed a precocious oocyte formation in CHX-treated heterozygotes (**Fig 5F**), further corroborating our RNA-seq and Ribo-seq findings.

To assess whether this regulatory shift extends to downstream targets of oocyte maturation, we analyzed the expression of *spn-4* and *meg-1*, which are regulated by OMA-1, OMA-2, and the TRIM-NHL RNA-binding protein LIN-41 (Tsukamoto et al. 2017; Huelgas-Morales and Greenstein 2018). Together, LIN-41 and the OMA proteins mediate a translational repression-to-activation switch critical for cytoplasmic oocyte maturation (Huelgas-Morales and Greenstein 2018). In CHX-treated heterozygotes, both *spn-4* and *meg-1* were overexpressed at the RNA level and displayed increased ribosome occupancy (**Fig 5G**). Notably, *lin-41* also showed increased mRNA abundance and ribosome occupancy in CHX-treated animals relative to controls (**Fig S6C, Table S5**). These results suggest that OMA proteins strongly promote *spn-4* and *meg-1* expression, potentially overriding LIN-41 mediated repression, and thereby drive premature oocyte maturation under elongation inhibited conditions.

Finally, titrating CHX in wild-type animals produced similar premature oocyte phenotypes at 250 µg/ml and 375 µg/ml (**Fig 5H**), further supporting the conclusion that moderate translation elongation inhibition alters the sperm-oocyte balance in *C. elegans*. Together, these results suggest how chronic ribosome stalling disrupts the normal sequence of gametogenesis, favoring oocyte pre-maturation and emphasizing the importance of controlled translation elongation in coordinating germline development.

## DISCUSSION

In this study, we identified the ribosomal protein RPL-36A as essential for cycloheximide (CHX) resistance in *C. elegans* and showed that the P55Q mutation maintains normal ribosome function under standard conditions while conferring resistance to this elongation inhibitor.

### CHX resistance as a selection marker for CRISPR co-conversion

Notably, heterozygous animals exhibited intermediate CHX resistance, clearly distinguishable from CHX sensitive animals. This property allowed us to use the *rpl-36A(P55Q)* mutation as a co-conversion marker for CRISPR-Cas9 editing, as an alternative to screening for roller or dumpy phenotypes (e.g., using *dpy-10(cn64)* or *sqt-1(e1350)* mutations) (Arribere et al. 2014). By leveraging CHX resistance to identify edited progeny, we improved the efficiency of identifying successfully edited candidates. Specifically, we obtained two lines of *rps-23(R67K)* among 30 candidates and one line of *rpl-16::HA::AVI::TEV* among 18 candidates. Despite this success, editing efficiency remained relatively low, likely reflecting the challenges of simultaneously modifying two highly conserved, essential and haploinsufficient ribosomal protein genes. In many cases, CRISPR-Cas9 mediated double stranded breaks in these loci may have been lethal, reducing the pool of viable co-edited lines.

Additionally, RPL-36A(P55Q) offers a convenient tool for positive selection in CRISPR genome editing without requiring large selection cassettes. Fluorescent tagging approaches that depend on antibiotic resistance (e.g., hygromycin) are quite efficient (Dickinson et al. 2015); however, this strategy may be unsuitable for essential genes, such as ribosomal proteins, that cannot tolerate large tags or display haploinsufficient phenotypes. In these cases, CHX-based selection is particularly advantageous for small edits or epitope tagging, significantly reducing the need for labor-intensive phenotypic screening, thus streamlining the isolation of genome edited strains.

### The absence of increased ribosome collisions and canonical stress responses

To understand how CHX affects ribosome dynamics, we performed high-resolution disome profiling (**Fig 3**). While most knowledge of ribosome stalling and stress responses originates from yeast (Simms et al. 2017, De and Mühlemann 2022), metazoans must integrate ribosome activity with complex developmental networks. In our heterozygous animals, carrying both CHX-sensitive and CHX-resistant ribosomes, none of these major stress response pathways (RQC, RSR, ISR) were detectably activated upon CHX treatment.

This lack of stress response suggests that chronic CHX-induced stalling differs from acute stalling events. One explanation is that the prolonged CHX exposure enables gradual adaptation, negating a sharp stress response. Indeed, disome profiling revealed fewer ribosome collisions, monosome analyses showed no major codon preferences. Moreover, markers of canonical stress responses, including RPS-10 ubiquitination (for RQC), eIF2α phosphorylation (ISR), PMK-1 activation, and transcriptional induction of ATF-4 targets, remained unaltered in CHX-treated heterozygotes.

We propose that CHX-sensitive ribosomes stall primarily at the start codon, reducing overall ribosome density along coding regions and thus increasing the effective distance between elongating ribosomes. This spacing lessens the likelihood of ribosome collisions and may help mitigate the impact of CHX by allowing the subset of initiating ribosomes to continue translating more efficiently.

### Impact of cycloheximide on germline development and oocyte maturation

Beyond its influence on ribosome dynamics, CHX treatment also altered germline development. Key regulators of cytoplasmic oocyte maturation, such as *spn-4* and *meg-1*, were overexpressed at the RNA level and showed increased ribosome occupancy under CHX. These findings align with the established roles of LIN-41 and OMA proteins in mediating translational control during meiotic progression (Huelgas-Morales and Greenstein 2018). Disrupting normal elongation thus leads to premature oocyte maturation, which is further supported by the upregulation of *fem-3* (**Table S5**), a key driver of the sperm-oocyte switch (Hodgkin 1986; Zanetti et al. 2012). Here, major sperm protein (MSP) signaling, which typically regulates oocyte maturation (Govindan et al. 2006), may be bypassed or downregulated, enabling oogenesis to occur without complete spermatogenesis. Intriguingly, this early oocyte maturation could theoretically promote outcrossing with males, potentially enhancing genetic diversity.

By generating heterozygotes with mixed CHX-resistant and CHX-sensitive ribosomes, our study establishes a *C. elegans* model to examine how partial translation elongation inhibition affects ribosome collisions, gene expression, and developmental plasticity. Taken together, our findings provide new insights into how chronic ribosome stalling can shape translational control metazoans and, in turn, redirect germline development by altering the balance between sperm and oocyte formation.

## ACKNOWLEDGEMENTS

We thank the members of the Sarinay Cenik Lab for their discussions and feedback. We also thank the Arribere Lab for sharing the strain WJA1025 and the Dickinson Lab for sharing the strain LP393. Some strains were provided by the CGC, which is funded by the NIH Office of Research Infrastructure Programs (P40 OD010440). This work was supported by the UT CNS Catalyst Grant, National Institutes of Health (NIH) NIGMS (R35GM138340), and Welch Foundation (F-2133-20230405) grants to E. Sarinay Cenik.

## AUTHOR CONTRIBUTIONS

Q.Z. and E.S.C. conceptualized the project and co-wrote the original manuscript. Q.Z. designed and conducted the experiments and performed the analyses. B.B. assisted with western blot experiments and manuscript editing. R.R. performed the motif enrichment analysis based on Ribo-seq and Diso-seq data and contributed to manuscript editing. R.T. and C.C. developed a Python-supported script to predict codons from Ribo-seq data. E.S.C. supervised the project and acquired funding.

## MATERIALS AND METHODS

### Generation of strains

Constructs, worm strains, and oligos sequences used in this study are listed in **Table S7, S8, and S9**.

The *rpl-36A(P55Q)* mutation allele was constructed using Cas9 protein under the control of *eft-3* promoter in pDD162, along with a gRNA targeting position P55 in the genomic sequence of *rpl-36A*. The gRNA was cloned into pQZ13, a derivative of pRB1017. The sgRNA construct pQZ13 was generated by the oligos ESC-QZ-92 and ESC-QZ-93. The homologous recombination template (ESC-QZ-96), which included the P55Q mutation, was synthesized by IDT. All plasmids for microinjection were purified using the Invitrogen PureLink HiPure Plasmid Miniprep Kit (#K210002).

The strain ESC217 (*cseIs2[rpl-36A(P55Q)] II*) was crossed to the balancer strain *mIn1* to generate the heterozygous strain ESC316 (*cseIs2[rpl-36A(P55Q)]/mIn1 II*). The strain ESC776 (*cseIs2[rpl-36A(P55Q)]/mIn1 II; cp145[mNG-C1^3xFlag::oma-2] V*) was generated by crossing the strain LP393(*cp145[mNG-C1^3xFlag::oma-2] V*) with the heterozygous strain ESC316.

### Worm growth

*C. elegans* strains were grown at 23 ℃ on agar plates containing nematode growth media (NGM) seeded with *Escherichia coli* strain HB101 for maintenance culture. To obtain synchronized embryos, adult animals were bleached using a buffer containing 0.5 N NaOH and 1.25% sodium hypochlorite for 6.5 minutes. Bleached embryos were placed onto NGM plates.

### Cycloheximide (CHX) treatment

The natural Cycloheximide (CHX) was purchased from Sigma (#01810). A 100 mg/ml stock solution in DMSO was prepared and stored at −20 ℃. CHX was diluted into the NGM agar and cooled to about 55 ℃ before pouring plates. Once solidified, plates were seeded with bacteria and left at room temperature for 1-2 days to allow bacterial lawn growth. Control plates for CHX experiments consisted of NGM plates with an equivalent concentration of DMSO. Additional CHX and control plates were prepared by spiking 500 µl water including CHX or DMSO onto pre-seeded plates.

### Sample and library preparation for RNA-seq, Ribo-seq and Diso-seq

Stage-matched L4 larvae with or without CHX treatment were collected in 50 mM NaCl and were cleaned from HB101 bacteria by sedimentation through a 5% sucrose cushion including 50 mM NaCl. After sucrose cleanup of bacteria, larvae were flash frozen in a worm lysis buffer (20 mM Tris-HCl (pH 7.4), 150 mM NaCl, 5 mM MgCl2, 1 mM DTT, 0.1% Triton X-100, 45 µM Emetine) and ground in liquid nitrogen with mortars and pestles. The frozen worm powder was thawed on ice, and 5 U/ml Turbo DNase (Thermo Fisher Scientific) was added. Then each lysate was divided into two aliquots.

For RNA-seq, around 1 ml TRIzol (Thermo Fisher Scientific) was added to the lysate, vortexed, and incubated for 5 minutes at room temperature. To extract RNA, 200 μl volume of chloroform was added and then the sample was mixed and spun at 15,000 rpm for 10 minutes. The aqueous layer was used for further RNA precipitation. Isolated RNA was isopropanol precipitated and 80% ethanol washed. Thermostable RNAseH (Lucigen) and a pool of 94 DNA oligonucleotides antisense to *C. elegans* ribosomal RNA were used to deplete rRNA from 100 ng total RNA (Arribere et al. 2016). RNA-seq libraries were prepared using SMARTer Stranded RNA-Seq kit (Clontech). Initially, RNA was alkaline fragmented at 95 ℃ for 4 minutes followed by the protocol optimized <10 ng RNA input. To amplify the sequences, 12 to 14 cycles of PCR were used. Library DNA was then purified using Agencourt AMPure XP beads (Beckman Coulter).

For Ribo-seq and Diso-seq, the lysate was vortexed briefly, and 5 U of RNAse I (Ambion) was added per ug of RNA, followed by incubation for 30 min at room temperature. The RNAse I reaction was stopped by addition of 25 mM ribonucleoside vanadyl complexes (Sigma). The lysate was loaded onto 34% sucrose made with the lysis buffer, and spun at 70,000 rpm using TLA 100.3 rotor (Beckman Coulter) for 4 hours at 4 ℃. The resulting pellet was solubilized in 1 ml of TRIzol to extract RNA. Resulting RNA was resolved on 10% Acrylamide TBE-urea gel (ThermoFisher Scientific), fragments between 26-34 nucleotides and 50-80 nucleotides were cut separately. The 26-34 nucleotide fragments were used for Ribo-seq, and the 50-80 nucleotide fragments were used for Diso-seq. Gel pieces were nutated in 30 mM Na-Acetate (pH 5.5) solution overnight in the cold room, with the eluate then precipitated as described above. Both Ribo-seq and Diso-seq libraries were generated using the D-Plex Small RNA-seq Kit (diagenode). The resulting libraries were quantified with Qubit dsDNA HS Assay Kit (ThermoFisher Scientific) and Agilent Bioanalyzer 2100. Libraries were sequenced on a NovaSeq X plus system to yield PE150 reads (Illumina), and sequenced on NovaSeq X Plus 10B flow cell (Illumina).

### RNA-seq, Ribo-seq and Diso-seq data analysis

Gene expression responses to CHX, derived from heterozygous animals carrying CHX-resistant and CHX-sensitive ribosomes, were analyzed at both the transcription and ribosome occupancy levels. For read mapping and further processing of the data, Riboflow nextflow (Ozadam et al. 2020) and edgeR pipeline (Robinson et al. 2010) were used (**Table S1, S2, S3, S4**).

For quantification and plotting of footprint density in metagene plots, nucleotide-mapped reads between 24–32 nt were used for monosomes, while a cutoff of 50–80 nt was applied for disomes (**Fig 3B–H**). To quantify the log fold change of monosome and disome occupancy in the metagene plots, position-specific changes were calculated in a matrix and normalized across conditions (**Fig 3D**) Monosome and disome occupancy (**Fig 3E, 3F, S2D, S4C**) were visualized using RiboGraph (Chacko et al. 2024).

To detect differences in translation efficiency (TE), ribosome-associated RNA (Ribo) levels, and overall RNA expression across the samples, three distinct contrasts were developed using edgeR. Specifically, the contrast for evaluating TE was defined as: TE_CHXvsControl = (CHX.Ribo – CHX.RNA) − (Control.Ribo – Control.RNA). These contrasts were then analyzed using a quasi likelihood F-test (glmQLFTest), with multiple testing corrections applied through a false discovery rate (FDR) method. An adjusted P-value threshold of 0.05 was used to determine significance (**Fig 5A, Table S5)**.

Gene Ontology (GO) enrichment analysis (Berriz et al. 2009) was separately performed on genes that were significantly over-or under-expressed at the RNA level or at the ribosome occupancy divided by RNA level, using Funcassociate 3.0 (Berriz et al. 2009). The GO categories with significant enrichments (p_adj_ < 0.05) were further filtered based on the following criteria: (1) a log odds ratio greater than 1, (2) fewer than 100 genes in the category, and (3) if two separate GO categories overlapped by more than 80%, one was randomly selected (**Fig 5C, S6A, S6B, Table S6**).

### Codon occupancy prediction

We developed a Python-supported script to predict codons based on Ribo-seq data (available on GitHub https://github.com/reikostachibana/ribopy_analysis).

### Motif enrichment analysis

In all replicates not treated with CHX, nucleotide sequences for disome and monosome reads were decoded using the mapping coordinates and reference genome. To obtain the amino acid sequence corresponding to these reads, the nucleotide sequences were translated with respect to the canonical start codon. Any portion of a read mapping outside the CDS or containing an incomplete codon was trimmed. For each amino acid and stop codon, the percentage of the total translated disome or monosome sequence was calculated. The same was performed for the coding sequence of all genes identified in disome and monosome reads. A random sample constituting 10% of all translated disome sequences was analyzed for motif enrichment utilizing STREME (Bailey 2021). For genes with disome reads containing the enriched motifs, coverage was plotted using Riboflow nextflow (Ozadam et al. 2020).

### Western blotting

Animals with or without CHX treatment were collected and cleaned from HB101 bacteria by sedimentation through a 5% sucrose cushion with 50 mM NaCl. The animals were then immediately flash-frozen in liquid nitrogen. Laemmli SDS sample buffer (Thermo Fisher, #AAJ61337AD) was added, and the samples were bead-beaten for 30 seconds before being boiled on a hot block for 5 minutes. Whole-worm lysates were separated on a 4%-12% Bis-Tris protein gel (Thermo Fisher Scientific, #NP0322BOX) and blotted onto a PVDF membrane.

Antibodies against Phospho-eIF2α (Cell Signaling Technology, #9721), Phospho-P38 antibody (Thermo Fisher Scientific, #MA5-15182), and Actin (MP Biomedicals, #8691001) were used at dilutions of 1:1000, 1:1000 and 1:500, respectively. HRP-conjugated secondary antibody (Cell Signaling Technology, #7074) and ECL reagents (Thermo Fisher Scientific, #34577) were used for detection.

To quantify western blots, TIFF images were captured for each blot using a chemidoc system, converted to 8-bit grayscale using Fiji, and the integrated intensity of each phospho-eIF2α, phospho-P38, and Actin band was calculated by Fiji. The phospho-eIF2α and phospho-P38 bands’ intensities were normalized by the corresponding Actin band intensity. Each normalized band intensity was expressed as a percentage of the control.

---

**Supplemental Figure 1.**
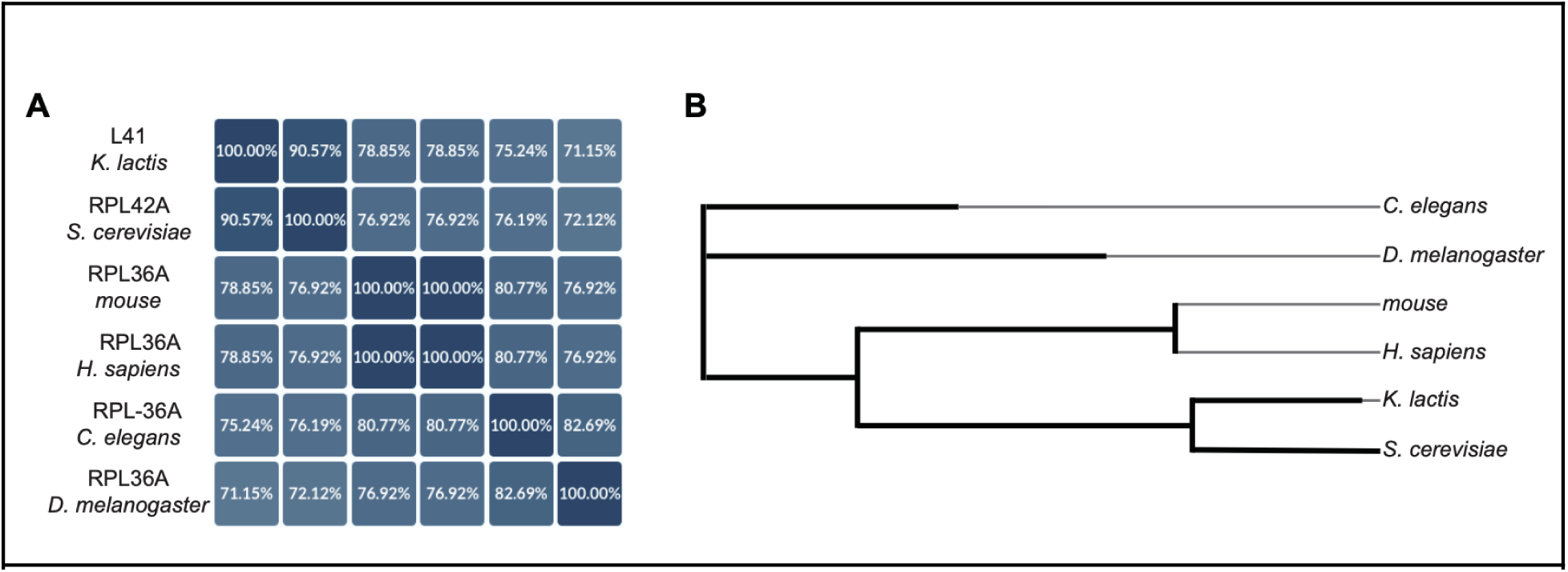
RPL-36A in *C. elegans* is homologous to *K. lactis* L41. **(A)** Comparison of the identity of RPL-36A in *C. elegans* with its homologs from *K. lactis*, *S. cerevisiae*, *mouse*, *H. sapiens*, and *D. melanogaster*. **(B)** Phylogenetic tree of *C. elegans* RPL-36A and its homologs.

**Supplemental Figure 2.**
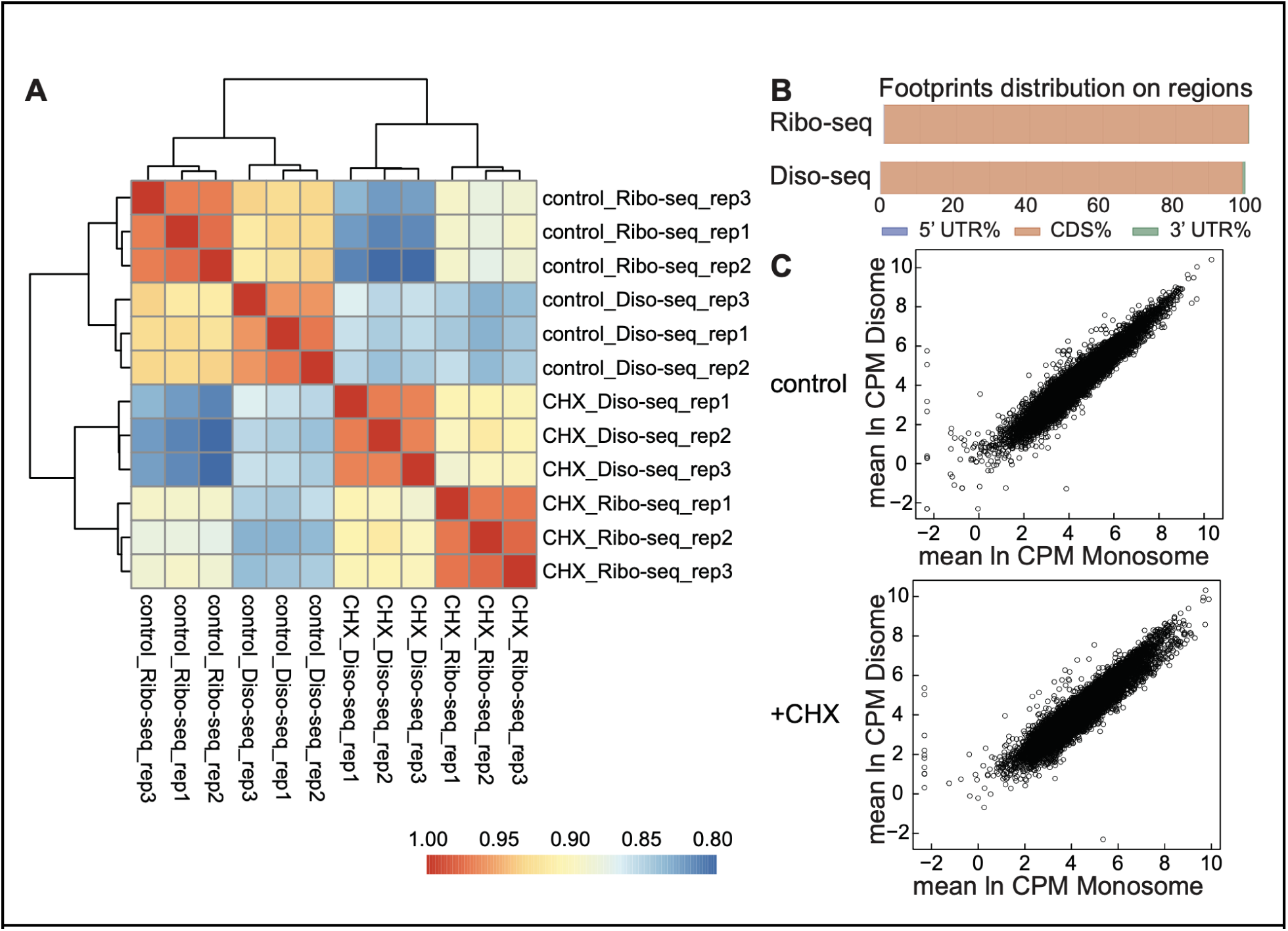
Data quality of Ribo-seq and diso-seq. (**A**) The heatmap of Pearson correlation coefficient of Ribo-seq and Diso-seq among all samples. **(B)** Reading frames of optimally mapped Ribo-seq reads and Diso-seq reads within annotated genes. **(C)** Scatter plots showing the correlation between disome abundance and monosome abundance in both CHX-treated animals and control samples. A higher frequency of ribosome collisions was observed in genes with higher monosome density in both conditions.

**Supplemental Figure 3.**
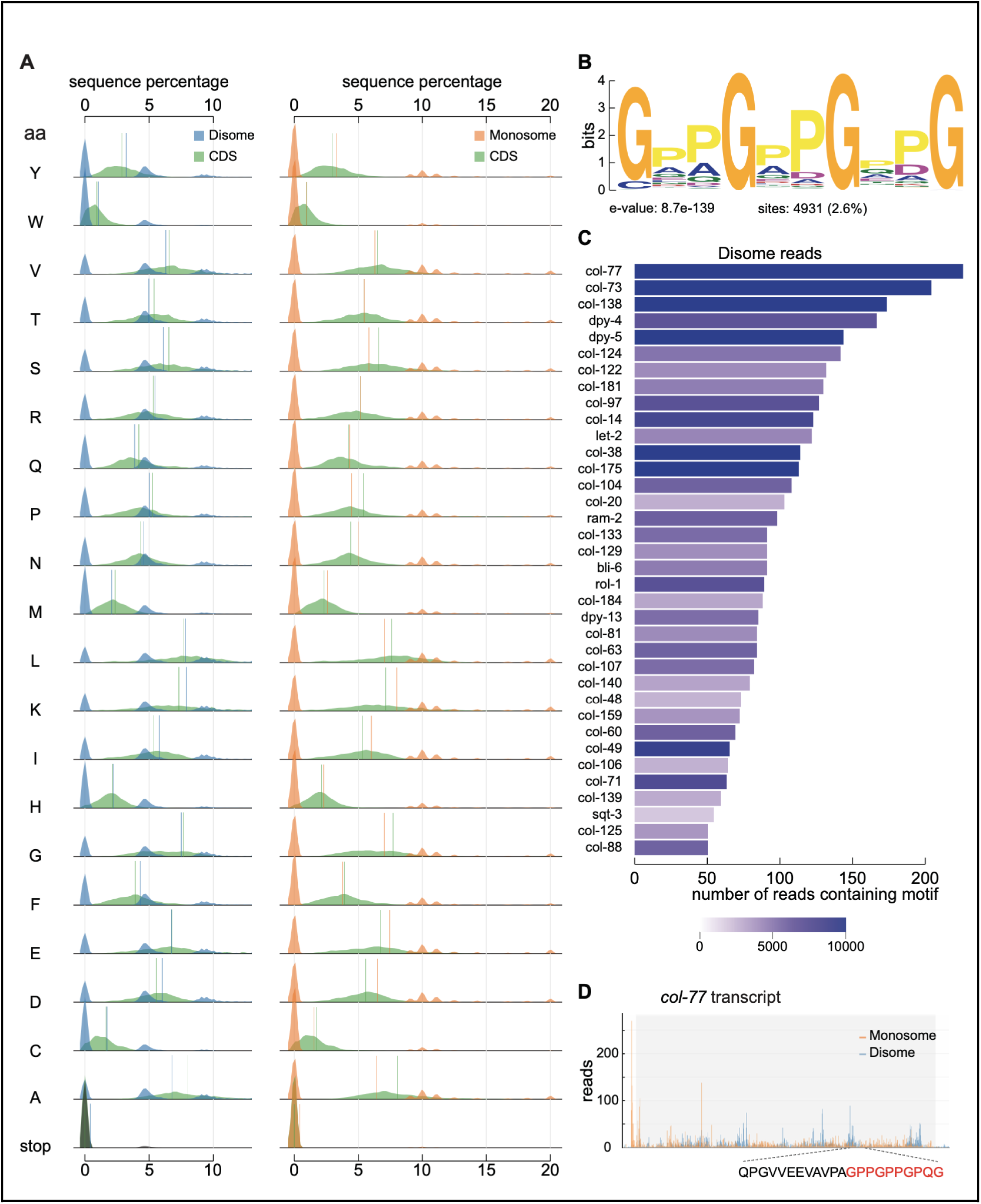
Relationship between ribosome stalling and motif enrichment. **(A)** Percentages of the 20 amino acids and stop codons within translated coding sequences (green), disome reads (blue), and monosome reads (orange), plotted as densities. Mean percentages are displayed as vertical lines for all three sequence categories. **(B)** Bit plot of the most highly enriched motif in translated disome reads (e-value = 8.7e-139, binomial test). **(C)** Bar chart depicting the number of disome reads within the STREME test set (containing 1% of total disome read sequences) which contain the motif in **(B)** for 36 genes. Total disome counts for each gene are demonstrated by the blue gradient, and only genes which have 50 or more reads containing the motif are shown. **(D)** Read coverage plot for *col-77*. The disome maximum containing the motif is labelled with the corresponding amino acid sequence and motif colored in red.

**Supplemental Figure 4.**
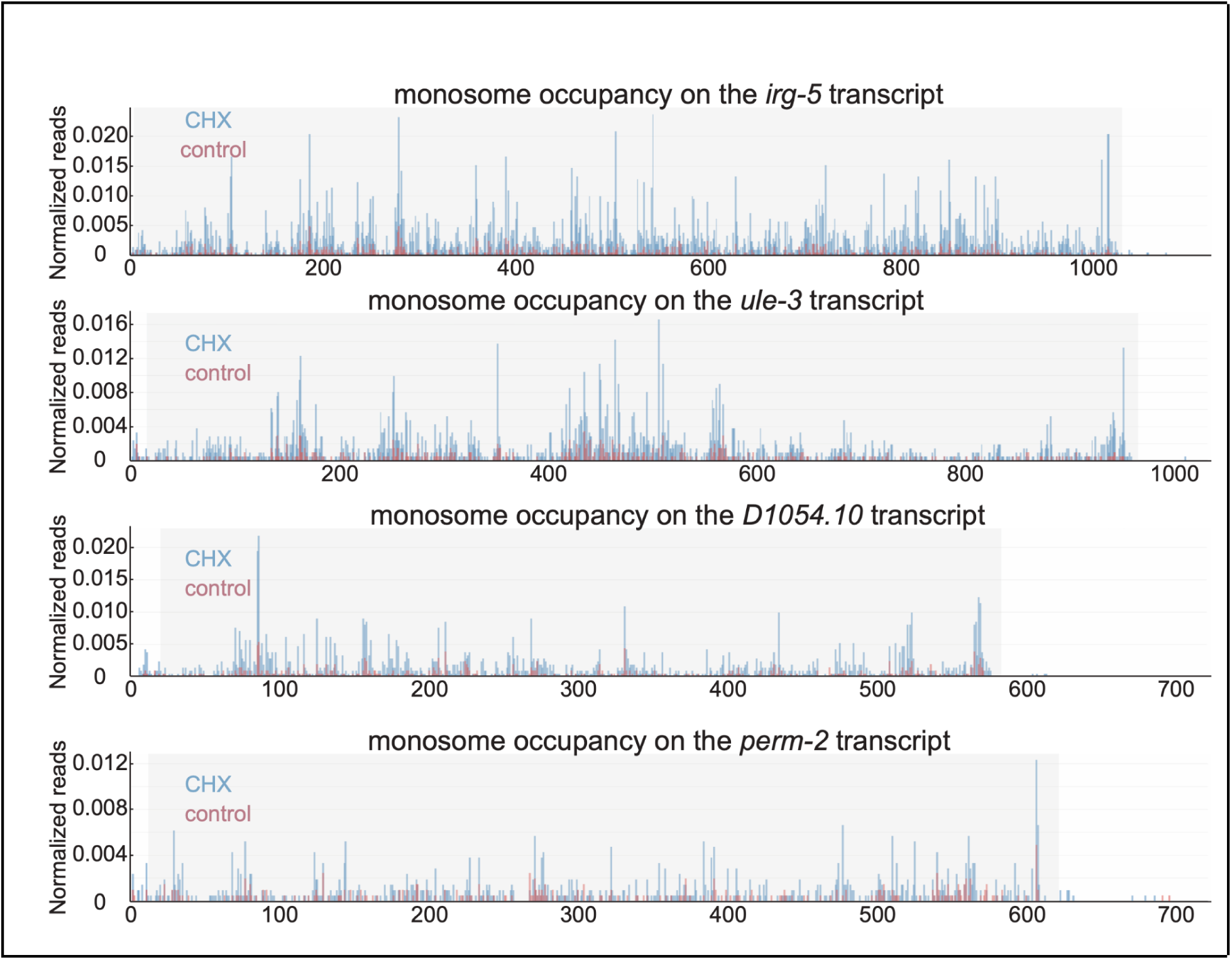
Distribution of monosome occupancy across gene transcripts. Representative examples of four genes (*irg-5*, *ule-3, D1054.10,* and *perm-2*) showing a significant increase in monosome occupancy per RNA in CHX-treated samples compared to control. The Y-axis represents the normalized value per 1,000 total reads. The light gray boxes along the X-axis indicate coding regions, while the white background represents the 5’UTR and 3’UTR, respectively.

**Supplementary Figure 5.**
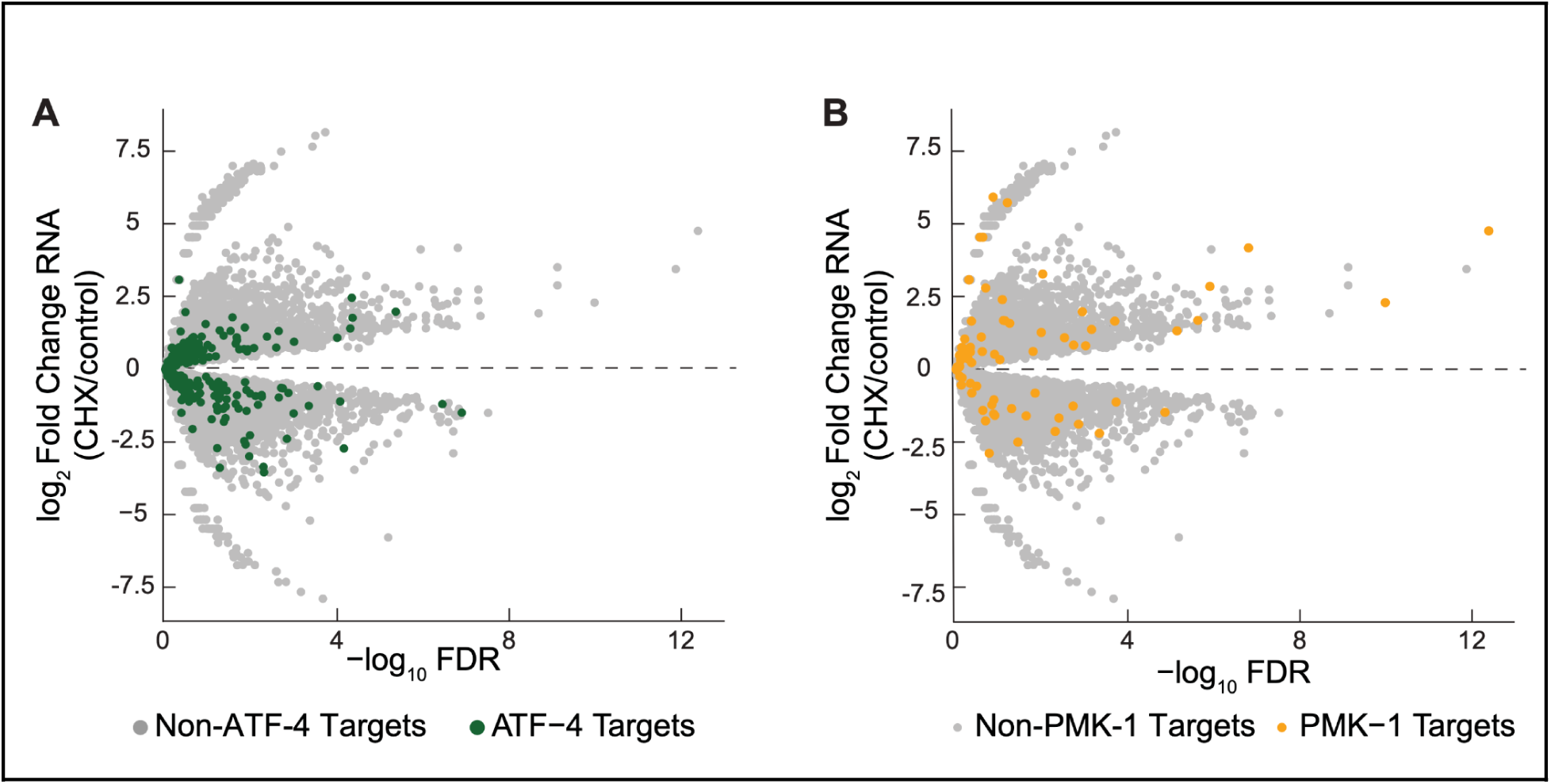
The expression signatures of ATF-4 and PMK-1 target genes in response to CHX treatment in heterozygous animals. **(A)** Log_2_ fold changes (CHX/control) of ATF-4 target genes (green) and non-target genes (grey) (y-axis) predicted by RNA-seq plotted against predicted -log_10_FDR (x-axis). **(B)** Log_2_ fold changes (CHX/control) of PMK-1 target genes (orange) and non-target genes (grey) (y-axis) predicted by RNA-seq plotted against predicted -log_10_FDR (x-axis).

**Supplementary Figure 6.**
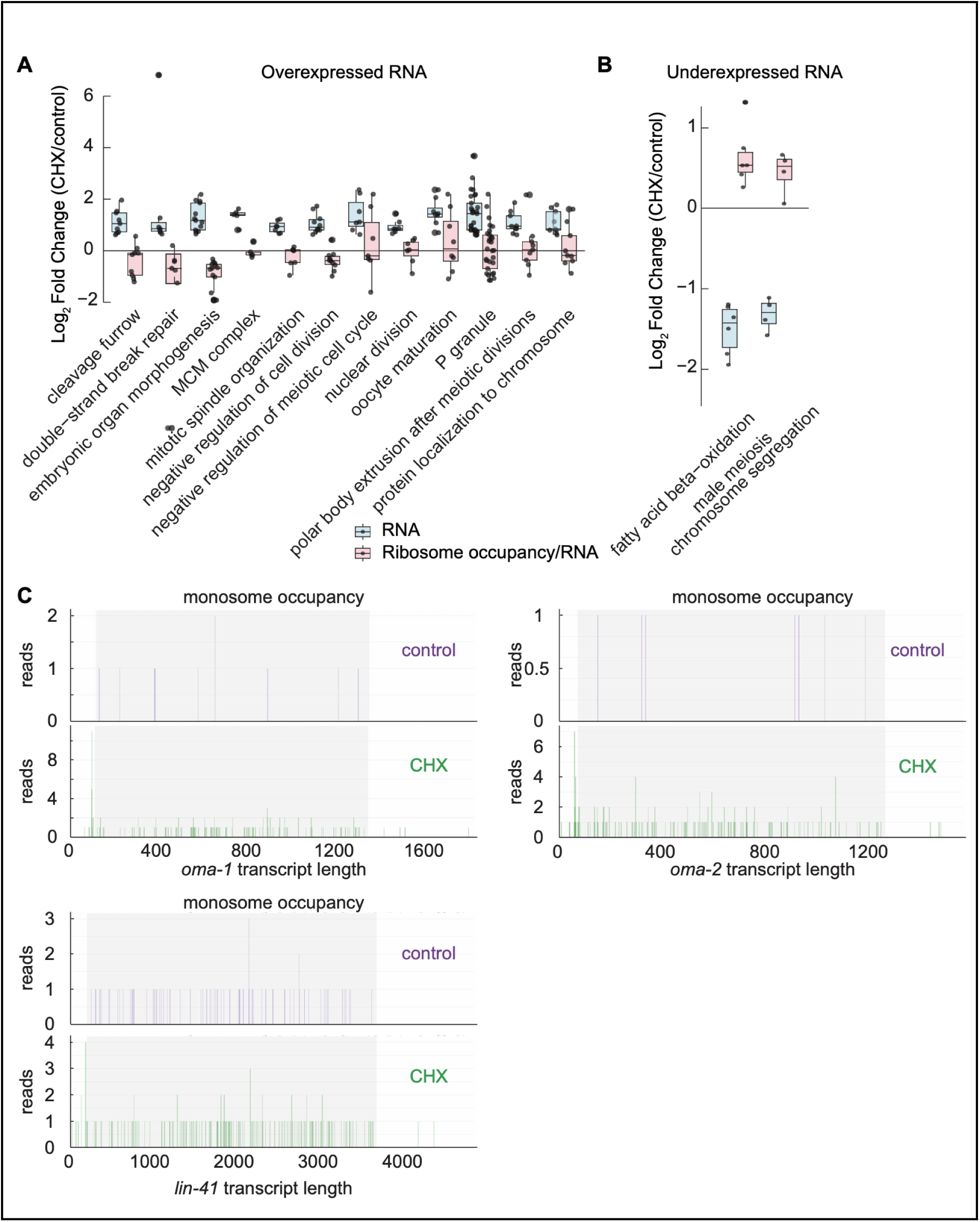
Significantly differentially expressed genes at the RNA level and ribosome occupancy in heterozygotes treated with CHX. **(A, B)** Log_2_ fold changes (CHX/control) in gene expression for representative GO categories enriched in RNA-overexpressed (**A**) and RNA-underexpressed (**B**) genes in heterozygous animals. The y-axis represents RNA-seq Log₂ fold-change values as determined by edgeR, with ribosome occupancy levels for these genes also plotted. A complete list of significantly enriched GO categories is provided in **Table S6**. (**C**) Monosome occupancy across *oma-1*, *oma-2*, and *lin-41* coding regions in CHX-treated and control animals. Monosome occupancy was enriched in CHX-treated animals for all three genes.

**Table S1. CPM-normalized RNA, monosome, and disome occupancy along transcripts in heterozygous animals with or without CHX xxtreatment.**

**Table S2. List of genes with a 2.7-fold higher disome occupancy than monosome occupancy and more than 100 reads each under normal conditions**

**Table S3. List of the all enriched motifs identified in disome-associated sequences.**

**Table S4. List of candidate genes with monosome occupancy profiles exhibiting a 2-fold increase in response to CHX treatment in heterozygous animals.**

**Table S5. Differentially expressed genes at RNA, ribosome occupancy, and ribosome occupancy/RNA levels in heterozygous animals in response to CHX treatment.**

**Table S6. GO enrichment analysis of genes with differentially expressed RNA and ribosome occupancy/RNA levels in heterozygous animals in response to CHX treatment.**

**Table S7. Constructs used in this study.**

**Table S8. *C. elegans* strains used in this study.**

**Table S9. Oligos used in this study.**

