## Supplemental Table 7, 8, 9 for "Cycloheximide resistant ribosomes reveal adaptive translation dynamics in *C. elegans*"

### **Table S6. Constructs used in this study.**

| **Constructs** | **Information** |
| --- | --- |
| pRB1017 | empty vector for gRNA cloning, Andrew Fire, Stanford University |
| pQZ13 | *rpl-36.A* sgRNA, generated from pRB1017 |
| pESC197 | *rps-23* sgRNA, generated from pRB1017 |
| pESC99 | *rpl-16 sgRNA*, generated from pRB1017 |
| PJA58 | *dpy-10 sgRNA,* Andrew Fire, Stanford University |
| pDD162 | *eft-3p::Cas9,* Andrew Fire, Stanford University |
| pC1(FD142) | Genomic *pha-1* gene, Andrew Fire, Stanford University |
| pQZ48 | *rpl-36.Ap::rpl-36.A(WT)::rpl-36.A 3'UTR* |
| pQZ49 | *rpl-36.Ap::rpl-36.A(P55Q)::rpl-36.A 3'UTR* |
| L3785 | *myo-3p::GFP,* Andrew Fire, Stanford University |
| pCFJ104 | *myo-3p::mCherry,* Andrew Fire, Stanford University |

### **Table S7 *C. elegans* strains used in this study.**

| **Strains** | **Genotype** | **Method** | **Source** |
| --- | --- | --- | --- |
| N2 | *WT* | NA | CGC |
| ESC217 | *cseIs2[rpl-36.A(P55Q)] II* | Microinjection | This study |
| ESC316 | *cseIs2[rpl-36.A(P55Q)]/mC6 II* | ESC217 cross with balancer strain *mC6* | This study |
| WJA1025 | [*rps-10*](https://cgc.umn.edu/strain/search?st1=rps-10&sf1=all)*(*[*srf1025*](https://cgc.umn.edu/strain/search?st1=srf1025&sf1=all)*[*[*rps-10*](https://cgc.umn.edu/strain/search?st1=rps-10&sf1=all)*::3*[*xHA*](https://cgc.umn.edu/strain/search?st1=xHA&sf1=all)*]) I* | NA | Arribere Lab, UCSC |
| LP393 | *cp145[mNG-C1^3xFlag::oma-2] V* | NA | Dickinson Lab, UT Austin |
| ESC776 | *cseIs2[rpl-36.A(P55Q)]/mC6 II; cp145[mNG-C1^3xFlag::oma-2] V* | ESC316 cross with LP393 | This study |

### **Table S8 Oligos used in this study.**

| **Oligos** | **Sequences** |
| --- | --- |
| ESC-QZ-92 | TCTTGTTCTTTCTGAAGATTGGCT |
| ESC-QZ-93 | AAACAGCCAATCTTCAGAAAGAAC |
| ESC-QZ-96 | TCCATTCTCAAGACGATTTTCTTGGTaGTCTTaGCCTTCTTTCTGAAGATTtGCTTaGTTTGTCCTCCGAATCCAGATTGTTTTCTGTCGTAACGACGACGTC |
| AF-ZF-827 | CACTTGAACTTCAATACGGCAAGATGAGAATGACTGGAAACCGTACCGCatGcGGTGCCTATGGTAGCGGAGCTTCACATGGCTTCAGACCAACAGCCTAT |
| ESC-QZ-182 | atgAGAATTGTCCTCGCTTGGGCCCCTGCCGAAGCCATCCCT |
| ESC-QZ-183 | ATAGGCGTATCACGAGGCCCTTAAGTTGAAAAGACGTTCTGATTATACCC |
| AF-ESC-377 | aattcataattttcagCGGTGTCGAAGCTAAGCAGCCTAACTCTGCTATCAAGAAGTGCGTTCGTGTCCAGCTCATCAAGAACGGAAAGAAGATCACCGCCTT |
| ESC-QZ-288 | GCCGCCCCAAAGATCGCTCAATACCAGAAGATCATCGAAGCCCTCGGATACAACGGAGGATCCGGATACCCATACGACGTCCCAGACTACGCCGAGAATCTGTACTTTCAATCCGGACTTAACGACATCTTCGAGGCCCAAAAGATCGAGTGGCACGAGTAAatccatgcacaaatctgttttgtttggtattataaaat |
